# Understanding the holobiont: crosstalk between gut microbiota and mitochondria during endurance

**DOI:** 10.1101/2021.01.08.425889

**Authors:** Núria Mach, Marco Moroldo, Andrea Rau, Jérôme Lecardonnel, Laurence Le Moyec, Céline Robert, Eric Barrey

## Abstract

Endurance exercise has a dramatic impact on the functionality of mitochondria and on the composition of the intestinal microbiome, but the mechanisms regulating the crosstalk between these two components are still largely unknown. Here, we sampled 20 elite horses before and after an endurance race and used blood transcriptome, blood metabolome and fecal microbiome to describe the gut-mitochondria crosstalk. A subset of mitochondria-related differentially expressed genes involved in pathways such as energy metabolism, oxidative stress and inflammation was discovered and then shown to be associated with butyrate-producing bacteria of the *Lachnospiraceae* family, especially *Eubacterium*. The mechanisms involved were not fully understood, but through the action of their metabolites likely acted on *PPARγ,* the *FRX-CREB* axis and their downstream targets to delay the onset of hypoglycemia, inflammation and extend running time. Our results also suggested that circulating free fatty acids may act not merely as fuel but drive mitochondrial inflammatory responses triggered by the translocation of gut bacterial polysaccharides following endurance. Targeting the gut-mitochondria axis therefore appears to be a potential strategy to enhance athletic performance.

## INTRODUCTION

To keep up with energy demand and to maintain homeostasis, endurance exercise modifies multiple systems, ranging from the whole-body level to the molecular level (Mach & Fuster-Botella, 2017; Clark & Mach, 2017). In recent years, our understanding of the role played by mitochondria during this kind of challenge has expanded far beyond its bioenergetic capacity, which is represented by well characterized pathways such as oxidative phosphorylation (OXPHOS), fatty acid β-oxidation (FAO) and the tricarboxylic acid (TCA) cycle (Pfanner *et al*, 2019). Indeed, it is now widely accepted that mitochondria regulate cytosolic calcium homeostasis and cellular redox status, that they generate much of the cell reactive oxygen species (ROS), and that they are involved in steroid and heme biosynthesis, urea degradation, apoptosis and initiation of inflammation through inflammasomes (Vezza *et al*, 2020; Jackson & Theiss, 2020; Wong *et al*, 2016; Chinnery & Hudson, 2013).

It has also been established that the mitochondrial genome (mtDNA) and the nuclear genome are constantly communicating with each other to regulate the aforementioned pathways. For example, most of the proteins involved in OXPHOS and mitochondrial functions like mtDNA replication and expression, mtDNA repair, redox and energy regulation are encoded by the nuclear genome and require specific targeting signals to be directed from the cytosol to mitochondrial surface receptors and then to the proper mitochondrial sub-compartments (Pfanner *et al*, 2019). The transcriptional programs of mitochondria comprise over 1,600 nuclear-encoded mitochondrial proteins (Pagliarini *et al*, 2008; Richter-Dennerlein *et al*, 2016). Any alteration in the OXPHOS and FAO processes, in mitochondrial membrane potential (ΔΨm), mitochondrial biogenesis, ROS production and inflammation can have a deep impact on the response to endurance exercise. For instance, increased mitochondrial biogenesis improves muscle endurance performance due to higher rates of OXPHOS and FAO (Hood *et al*, 2011). On the contrary, lower metabolic rates, increased ROS production and acidosis during prolonged exercise are associated with fatigue and inability to maintain speed (Rapoport, 2010). Endurance or long exercise and mitochondrial functions are strictly intertwined and influence each other.

The gut microbiota is considered a central organ because of its direct and indirect roles in host physiology, including improved metabolic health and athletic performance (Keohane *et al*, 2019; Barton *et al*, 2017; Scheiman *et al*, 2019). A healthy ecosystem in horse includes numerous, highly dominant taxa along with a multitude of minor players with lower representation, but important metabolic activity (Mach *et al*, 2020; Costa & Weese, 2018; Kauter *et al*, 2019).

The interdependence of gut microbiota and mitochondria is being increasingly recognized, with many diseases originating from mitochondrial dysfunctions linked to well-described changes in gut microbiota (Cardoso & Empadinhas, 2018; Bajpai *et al*, 2018; Karlsson *et al*, 2013; Yardeni *et al*, 2019; Franco-Obregón & Gilbert, 2017; Saint-Georges-Chaumet *et al*, 2015; Mottawea *et al*, 2016; Saint-Georges-Chaumet & Edeas, 2018). This complex interplay occurs principally through endocrine, immune and humoral signalling (Mottawea *et al*, 2016). Enteric short-chain fatty acids (SCFAs), the major class of metabolites produced from bacterial fermentation of non-digestible carbohydrates, are widely thought to mediate the relationship between the gut microbiota and the mitochondria in different tissues. Branched-chain amino acids (BCAAs), secondary bile acids, ROS, nitric oxide (NO), and hydrogen sulfide (H_2_S) are also thought to play at least a partial role in this molecular interchange (Clark & Mach, 2017).

While formal proof is missing in horse athletes, a prevailing hypothesis is that the gut microbiota and its metabolites regulate crucial transcription factors and coactivators involved in mitochondrial functions that underpin endurance performance (Hawley *et al*, 2018). In mice models, gut microbiota depletion via broad-spectrum antibiotics showed reduced production of SCFAs, lower bioavailability of serum glucose, decreased endurance capacity and impairment of the *ex vivo* skeletal muscle contractile function (Nay *et al*, 2019). In close agreement, gut microbiota depletion also triggered a reduction of both faecal SCFA content and circulating concentration of SCFAs coupled to a drop in running capacity in mice (Okamoto *et al*, 2019). In contrast, mice with *Veillonella* in their intestinal ecosystem showed significantly increased submaximal treadmill run time to exhaustion (Scheiman *et al*, 2019), prompting the authors to speculate that the lactate generated during sustained bouts of exercise could be accessible to the microbiota and converted into SCFAs that ultimately enhanced energetic resilience and stamina. Alternatively, there may be other mechanisms through which gut microbiota and its metabolites relate to mitochondria, including but not limited to the regulation of mitochondrial oxidative stress (Jones & Neish, 2014; Franco-Obregón & Gilbert, 2017), as well as the activation of the inflammasome and the production of inflammatory cytokines, all of which are key players in the adaptation to endurance exercise (Clark & Mach, 2017; Mach & Fuster-Botella, 2017).

We recently presented a three-pronged association study, connecting horse gut microbiota with untargeted serum metabolome data and measures of host physiology and performance in the context of endurance (Plancade *et al*, 2019). We found no significant associations between the gut ecosystem and serum metabolites, especially those relying heavily on mitochondrial OXPHOS, FAO, TCA and gluconeogenesis. The number of annotated metabolites in the study was likely insufficient to reliably encompass the given mitochondrial functions. To further advance our knowledge of the molecular basis for the gut-mitochondria crosstalk that support the adaptation to long exercise, in this work we tethered whole blood transcriptome profiling to our previous metabolome and metagenome data. In doing so, we sought to identify the ways in which mitochondrial and nuclear transcriptomes coordinate with each other, and how gut microbiota and circulating metabolites can dynamically modulate this process. By jointly characterizing the whole blood transcriptome, metabolome, fecal microbiota and SCFAs of 20 elite horses competing in an endurance race, we aim to provide a functional readout of microbial activity and improve our understanding of the gut microbiota-mitochondria axis during long exercise.

## RESULTS

To elucidate how mitochondria and gut microbiota are linked during endurance exercise, we studied 20 healthy endurance horses selected from the cohort already described by Plancade *et al*. (2019) and Le Moyec *et al*. (2019) (Fig 1). All of the animals were of similar age and performance level (Table EV1). They performed a long exercise during about 8 hours at an average speed of 17.1 ± 1.67 km/h with some rest periods every 30-40 km.

**Figure 1.**
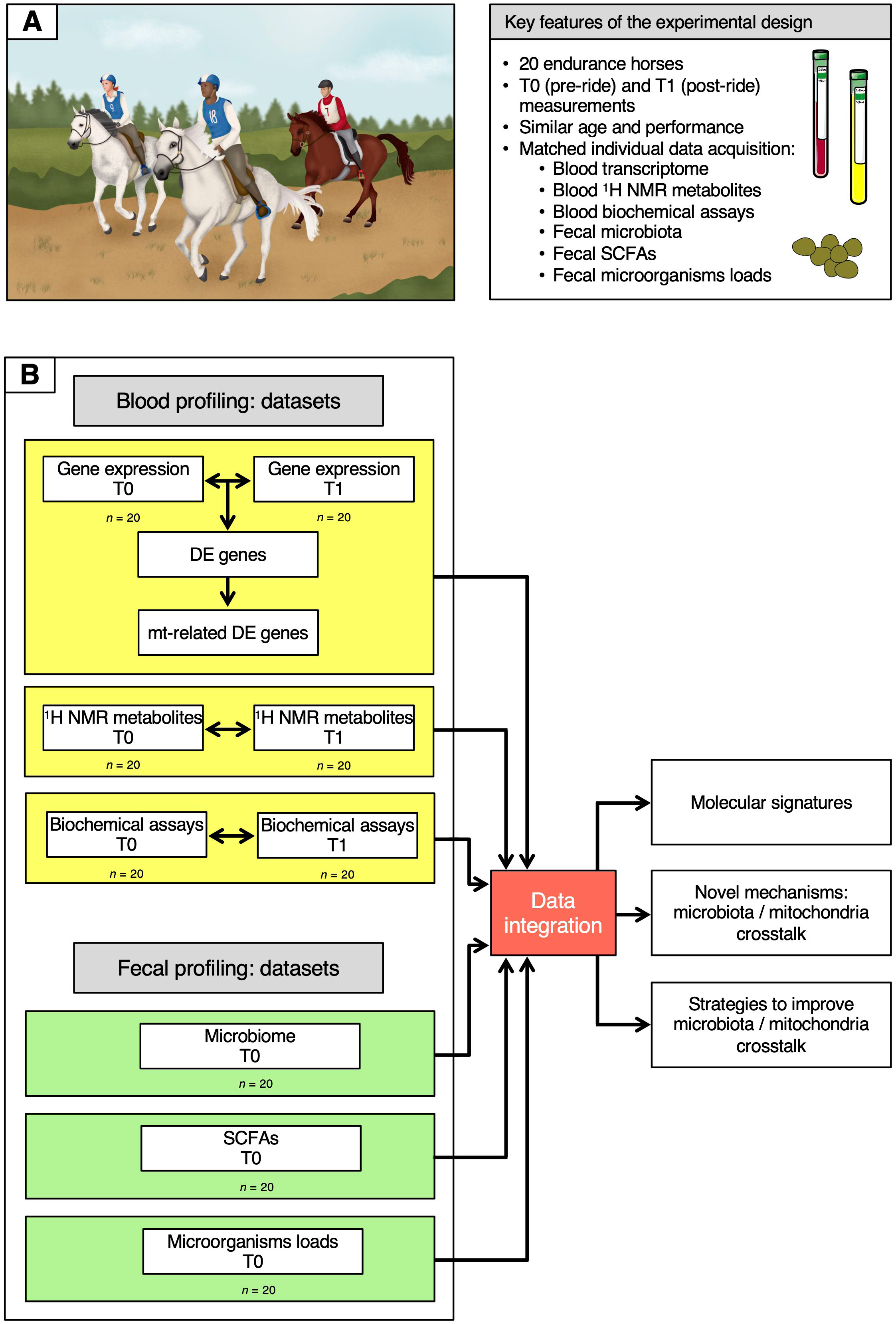
Description of the cohort and of the data analysis workflow. A Key features of the experimental design. B Overview of the data analysis workflow. On the left, the six datasets used in the study are depicted, indicating whether they were obtained at T0 and T1 or at T0 only. Data information: written permission for publication of the drawings corresponding to endurance event in the panel A was obtained. In the A panel, the pictures of the blood and plasma tubes were download from In all cases, no changes were made. Servier Medical Art by <u>Servier</u> is licensed under a <u>Creative Commons Attribution 3.0 Unported License.</u>

Whole transcriptome profiles, proton nuclear magnetic resonance (^1^H NMR) metabolome profiles and biochemical assay data were obtained from blood samples collected at both T0 (pre-ride) and T1 (post-ride), while SCFAs measurements, 16S rRNA data and the concentration of bacteria, anaerobic fungi and ciliate protozoa were generated from fecal samples at T0 alone.

### Blood transcriptome profiles and mitochondrial related genes

To gather information about the global structure of the blood transcriptome, a scaled principal component analysis (PCA) was carried out on the expressed genes (*n* =11,232). The first component accounted for 43% of the total variability and revealed a marked separation of the two time points (Fig 2A). We then carried out a standard differential analysis between the two time points. After Bonferroni correction of the raw *p*-values, a total of 6,021 differentially expressed genes (DEGs) was obtained at an adjusted *p* < 0.05, of which 2,658 were upregulated and 3,363 downregulated at T1 respect to T0 (Table EV2; Fig 2B).

**Figure 2.**
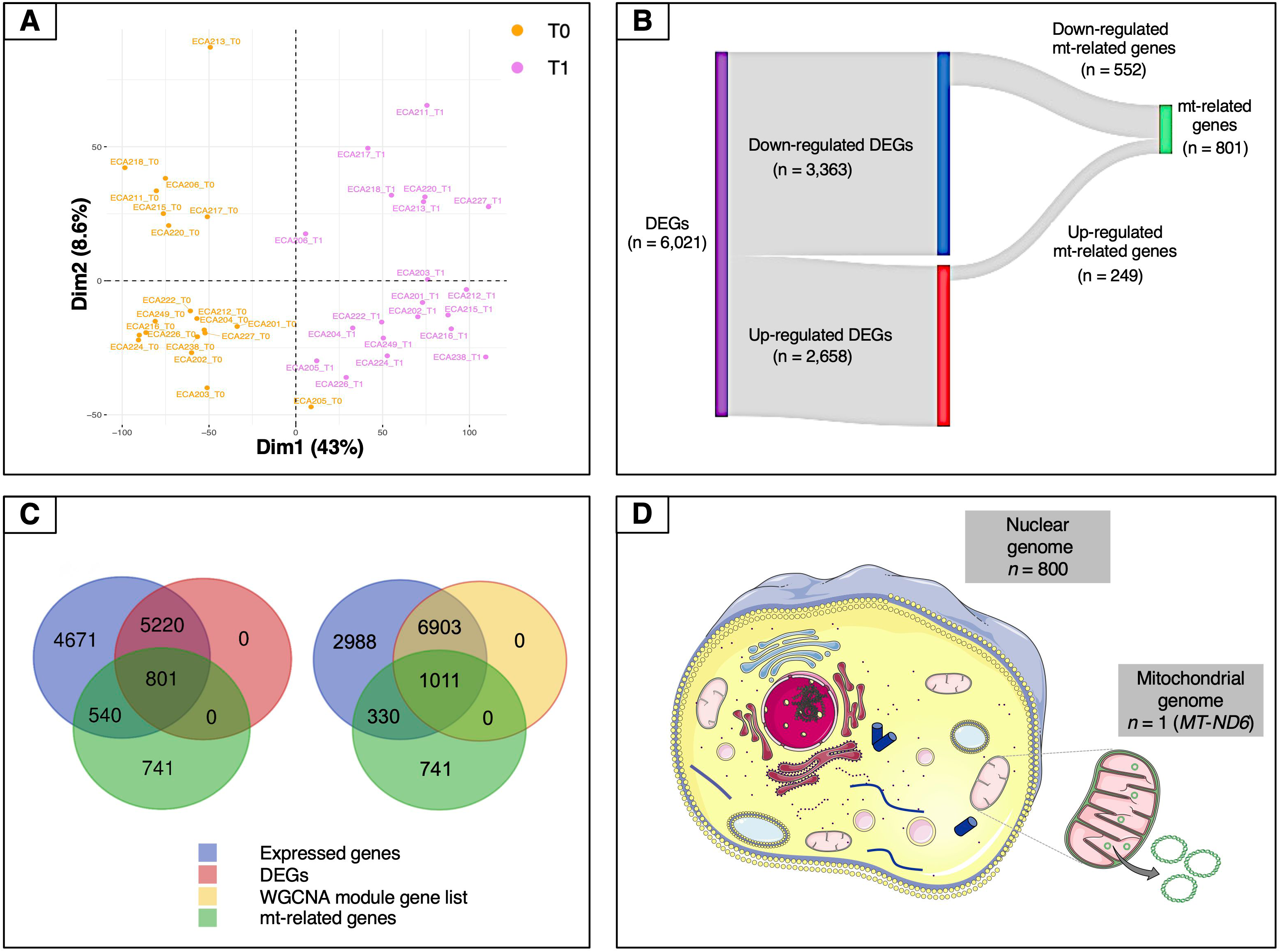
Overview of the mt-related genes. A Plot of the first two components of the PCA obtained using all of the expressed genes. B Sankey diagram of the differentially expressed genes, showing the numbers of up-regulated, down-regulated and mt-related genes. Only a relatively small fraction of the mt-related genes (249 out of 801) was up-regulated, whereas most (552 out of 801) were down-regulated. C Venn diagrams illustrating the overlaps among expressed genes (blue), differentially expressed genes (purple), genes included in WGCNA modules (yellow) and mt-related genes (green). D Illustration of the proportion of mt-related genes encoded by the mitochondrial and by the nuclear genomes. Except for *MT-ND6*, all of the other genes are encoded by the nuclear genome. Data information: in panel D, the picture of the mitochondria was download from In all cases, no changes were made. Servier Medical Art by Servier is licensed under a <u>Creative Commons Attribution 3.0 Unported License.</u>

These results were then complemented using a weighted gene co-expression network analysis (WGCNA) (Langfelder & Horvath, 2008) on the expressed genes. WGCNA identified three gene modules, corresponding to 7,914 genes, that were correlated to the ^1^H NMR and biochemical assay metabolites (Table EV3). These genes strongly overlapped with the set of DEGs, 91.1% of which (i.e., 5,486 out of 6,021 genes) were included among them. The metabolites which showed the highest levels of correlation with the gene modules were bilirubin, non-esterified fatty acids (NEFAs), tyrosine, lactate and, to a lesser extent, β-hydroxybutyrate (BHB) (see next paragraph, Fig EV1).

Because we were especially interested in understanding the role played by mitochondria in our biological system, we then decided to study in more detail the features related to these organelles. To this end, we used a literature-based meta-analytic approach to build a non-redundant consensus list of 2,082 genes related to mitochondria, based on the information available in the Integrated Mitochondrial Protein Index (IMPI) (Smith & Robinson, 2019), the Mitocarta Inventory (Calvo *et al*, 2016) and the literature (Expanded View Information and Table EV4). This consensus list was also descriptively annotated to gain global insight into the main biological functions represented within it. A total of 80 third-level KEGG hierarchies were identified, with a strong representation of pathways related to carbohydrate and lipid metabolism (8 and 9 ontology terms, respectively). Eight pathways were associated with amino acid metabolism, while five were linked to the apoptosis process (Fig EV2). This subset was crossed with each of the DEG and WGCNA module gene lists. In the case of the DEG list, the intersection with this subset yielded a total of 801 genes (Table EV5 and Fig 2C). Both the Fisher’s exact test and the hypergeometric test showed strong levels of over-representation, with *p*-values of 1.0 x 10^-5^ and 9.02 x 10^-7^, respectively. In the case of the WGCNA gene modules, the intersection included 1,011 genes (Fig 2C). Again, both statistical tests indicated strong enrichment, with *p*-values of 1.0 x 10^-5^ and 1.07 x 10^-5^, respectively.

The set of 801 genes in the intersection of the mitochondrial consensus and DEG list, which will be referred to hereafter simply as “mt-related genes”, was selected for the downstream steps of analysis. All of the mt-related genes were encoded by the nuclear genome except for *MT-ND6* (MT-NADH dehydrogenase, subunit 6), which was encoded by the antisense strand of the mt-DNA (Fig 2D). These mt-related genes were further characterized to gather information about their molecular function. The functional analysis showed that roughly 75 % of these genes were directly involved in energy metabolism (i.e., pathways such as OXPHOS and FAO) and metabolite synthesis and degradation (Fig EV3). For instance, we observed an enrichment of genes related to nutrient transport across the mitochondrial inner membrane (*TOM/TIM* units, *VDAC*, *MPC1*, *ACAA1*, members of the mitochondrial carrier family *SLC25* and of the pyruvate dehydrogenase kinase isozyme), fatty acid metabolism (*ELOVL7, SIRT5,* and *ACAD* members), lipogenesis *(FASN* and *PPARγ),* and fatty acid channelling into oxidation (*CPT1B, ACADVL, ACOT9, ACOX1, ACSL1, ACSL4, ACSS3*). Additionally, key genes involved in the mitochondrial biogenesis (*POLG* and *POLG2*), mitochondrial fission (*PINK1*), mitochondrial fusion (*OPA1*), mitophagy (*BNIP3* and *PINK1),* oxidative stress (*SOD1, SOD2,* glutathione S-transferases and glutathione peroxidases families), and the resolution of lipopolysaccharide induced pro-inflammatory pathway (*C1QBP*) were also found in this list. *CREB*, the most potent activator of *PGC-1α* (Wu *et al*, 2006) was also differentially expressed upon endurance exercise.

Our data further indicated that among the mt-related genes, at least 21 genes encoding rps and rpl proteins of the small and large subunits of ribosomes (*rpl3, rpl4, rpl5, rpl6, rpl18, rpl23, rpl27, rpl36, rps3, rps8, rps9, rps10, rps11, rps12, rps13, rps14*, *rps15, rps16, rps17, rps18, rps19*) (Janouškovec *et al*, 2017; Esser *et al*, 2004; Maier *et al*, 2013) were common with the α-Proteobacteria. This was also the case for the methionine sulfoxide reductase A (*MSRA*) and NAD(P)H dehydrogenase quinone 1 (*NQO1*) (Crisp *et al*, 2015), consistent with a vertical origin in the mitochondrial endosymbiont.

### 1H NMR metabolome, biochemical assay and acetylcarnitine profiles

The ^1^H NMR metabolome and biochemical assay profiles used in this paper were gleaned from our previous works (Plancade *et al*, 2019; Le Moyec *et al*, 2019). Briefly, a total of 50 ^1^H NMR known metabolites was detected in the plasma, including several amino acids, energy metabolism-related metabolites and organic osmolytes (Table EV6). Three well-known microbial derived metabolites were ascertained, including formate, dimethyl sulfone and trimethyl amine oxide (TMAO).

The relative abundance values of these circulating metabolites fell within the normal reference range for healthy horses. However, the concentration of lactate (a proxy for glycolytic stress and disturbances in cellular homeostasis (Hawley *et al*, 2018)) was significantly increased after the race, as well as the levels of fatty acids from lipoproteins and of certain amino acids, namely alanine, branched amino acids such as leucine, valine and *iso*-valerate, glutamate, glutamine and aromatic amino acids such as tyrosine and phenylalanine. Ketone bodies were slightly increased after the race (i.e., acetoacetate and acetate; Table EV6). In the case of biochemical profiles, all of the horses showed above-average concentrations for total bilirubin, creatine kinase, aspartate transaminase, and serum amyloid A after the race. The NEFA and BHB concentrations also showed similar patterns (Table EV7).

### Fecal short chain fatty acids measurements, 16S rRNA data and microorganism concentrations

The microbiota composition and derived-metabolites were obtained from Plancade *et al*. (2019), but it is important to note that the 16S rRNA raw sequences were re-analysed using the QIIME 2 plugin, which quantitatively improved results over QIIME 1 by enhancing the pre-processing of sequenced reads, the taxonomy assignment, the phylogenetic insertion and the generation of amplicon sequence variants (ASVs) (Callahan *et al*, 2017). A total of 519,866 high-quality sequence reads were obtained (mean per subject: 21,131 ± 15,625, range: 6,036 – 57,389). Reads were clustered into 3,384 chimera- and singleton-filtered ASVs at 99% sequence similarity (Table EV8). The intestinal microbial community found in the total set of 20 individuals as a whole was made up of a core of 23 genera, the core being defined as the genera shared by 99% of all sampling events with a minimum 0.1% mean relative abundance. Overall, 61% of the core genera belonged to the Firmicutes phylum, mainly to the *Lachnospiraceae* and *Ruminococcaceae* families (Fig EV4A). A total of 100 unique genera was identified in the microbiota (Table EV9). The majority of these genera (80%) fell among the 20 most abundant (Fig EV4B) and accounted for more than 75% of the sequences in the data (Table EV9). To deeper understand how the gut microbiota functioned, SCFAs, pH measurements and the loads of anaerobic fungi, protozoa and total bacteria in feces were also investigated. The main products of microbial fermentation were acetate, propionate and butyrate, with ratios ranging from 60:32:8 to 76:19:5. Small amounts of branched chain fatty acids (*iso*-butyrate, valerate and *iso*-valerate) were also detected (Table EV10). Although bacteria represented the major portion of the fecal microbiota in our horses, the relative concentrations of anaerobic fungi and ciliate protozoa were 0.82 and 0.76, respectively (Table EV11).

### Integration of transcriptome, ^1^H NMR metabolites, biochemical parameters, fecal microbiota, SCFA and microorganism concentrations

After the identification of the 801 mt-related genes that were regulated by endurance exercise, it remained to be determined which genes were interconnected to the gut microbiota and responded to specific circulating molecules. To this aim, we applied four independent statistical methods using the mt-related genes as the response variable and the other data sets, namely ^1^H NMR metabolome, biochemical assay profiles, fecal microbiota, fecal SCFAs and the concentrations of bacteria, anaerobic fungi and protozoa as exploratory variables.

We first used global non-metric multidimensional scaling (NMDS) ordinations to visualize the structure of mt-related gene expression (ordination stress = 6%, *k* = 2, non-metric fit *r*^2^ = 0. 0.996, linear fit *r*^2^ = 0.988) and we then fitted all sets of explanatory variables to the ordination to find the most influential variables (Fig 3A). Bacteria such as *Oribacterium, Rikenellaceae* RC9, *Ruminococcaceae* NK4A214, unclassified rumen bacterium and *Clostridium sensu stricto* showed the strongest correlation to all ordinations, together with some microbial-derived metabolites (i.e., dimethyl sulfone, formate, valerate and *iso*-valerate) and the plasmatic NEFA (adjusted *p* < 0.05; Fig 3A-B).

**Figure 3.**
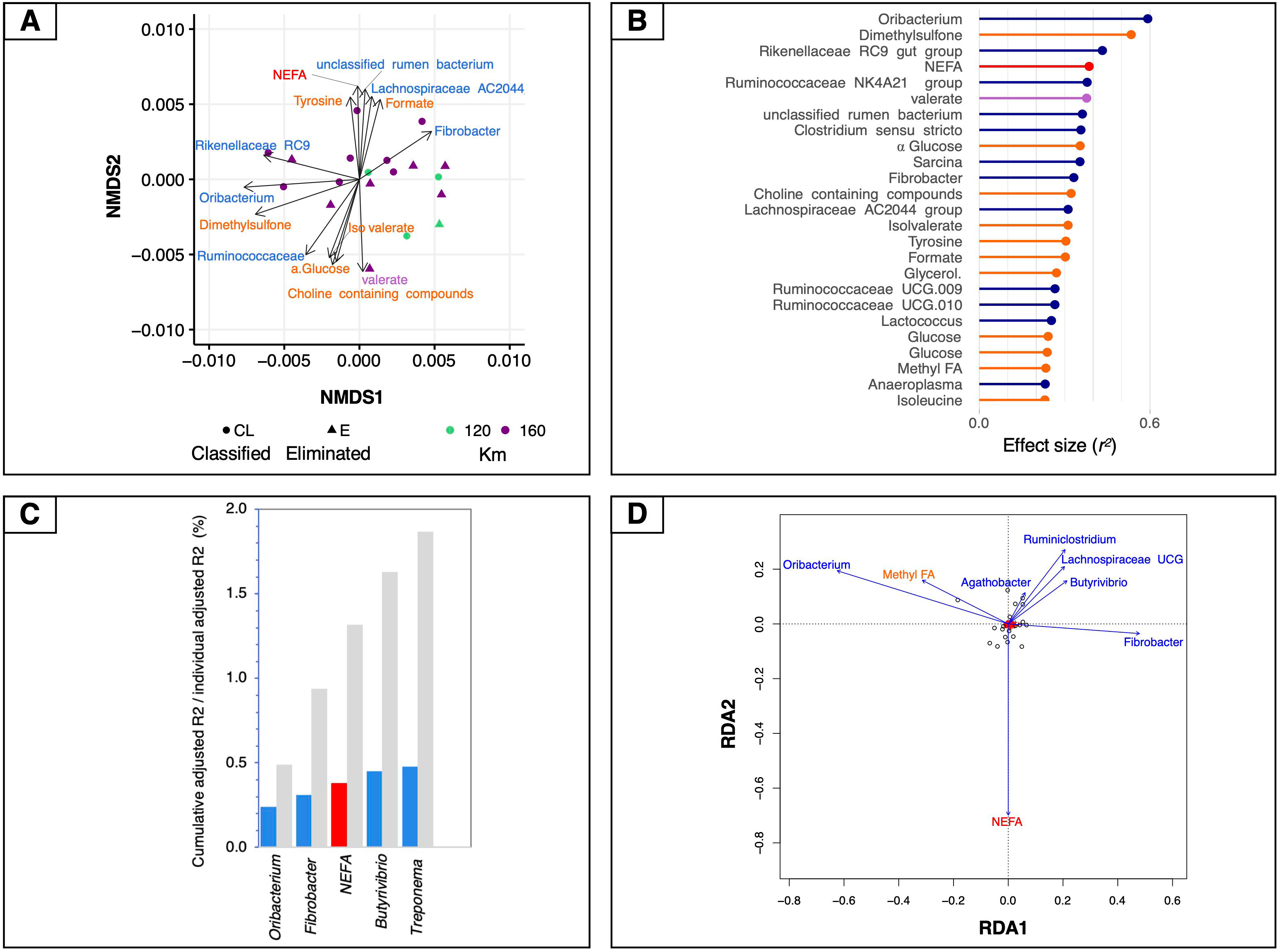
Associations between mt-related genes, microbiota and circulating metabolites. A Dissimilarities in mt-related gene expression represented by the non-metric multidimensional scaling (NMDS) ordination plot. The Bray–Curtis dissimilarity index was calculated on normalised data, the samples were coloured according to the total length of the race and the two different shapes of the dots indicate if the horses finished the race or if they were eliminated. B Effect sizes of gut microbiota, fecal SCFAs, ^1^H NMR and biochemical assay metabolites over NMDS ordination. Covariates are coloured according to the type of dataset: ^1^H NMR metabolites are in orange, biochemical assay metabolites in red, fecal SCFAs in violet and bacteria in dark blue. Horizontal bars show the amount of variance (*r*^2^) explained by each covariate in the model as determined by ‘envfit’ function. C Grouped bar chart showing the cumulative effect sizes of covariates on mt-related gene expression (coloured bars) compared to individual effect sizes assuming covariate independence (grey bars) using a stepwise model selection using distance-based redundancy analysis (dbRDA). Covariates are coloured according to the type of dataset: red for biochemical assay metabolites, and blue for bacteria. D Plot showing the covariates that contribute significantly to the variation of mt-related genes determined by stepwise model selection using redundancy analysis (RDA). The arrows for each variable show the direction of the effect and are scaled by the unconditioned *r*^2^ value. Covariates are coloured according to the type of dataset: red for biochemical assay metabolites and dark blue for bacteria.

To control for spatial variance and to identify the minimal combination of non-redundant covariates that would best fit with the mt-related gene profiles, a more rigorous multivariate distance-based redundancy analysis (db-RDA) was used on constrained NMDS ordinations. In agreement with the aforementioned results, the expression of mt-related genes responded most strongly to bacteria such as *Treponema* (*r*^2^_adj_ = 0.48, *p* = 0.007), followed by *Butyrivibrio* (*r*^2^_adj_ = 0.45, *p* = 0.002), plasmatic NEFA (*r*^2^_adj_ = 0.38, *p* = 0.003), *Fibrobacter* (*r*^2^_adj_ = 0.31, *p* = 0.0008) and *Oribacterium* (*r*^2^_adj_ = 0.24, *p* = 0.003; Fig 3C). The ^1^H NMR metabolites and fecal SCFAs and the concentration of fecal microorganism did not directly contribute to the variation of mt-related genes. Therefore, they were not selected by the dbRDA model.

These findings were further confirmed by an RDA forward-selection model based on the Akaike information criterion. Specifically, *Oribacterium* (*F* = 7.26, *p* < 0.005)*, Fibrobacter* (*F* = 2.96, *p* < 0.005), *Butyrivibrio* (*F* = 2.91, *p* < 0.005), *Agathobacter* (*F* = 2.12, *p* < 0.005), *Treponema* (F = 7.15, *p* < 0.01), unclassified rumen bacterium (*F* = 1.96, *p* < 0.01), and the concentration of plasmatic NEFA (*F* = 2.91, *p* < 0.005) explained most of the variance observed in mt-related genes (Fig 3D). However, the ^1^H NMR metabolites, the fecal SCFAs and concentration of microorganisms were not found to contribute significantly to the variability in mt-related gene expression. The first constrained axis (RDA1) explained 44% of the variance in mt-related gene expression, and the second (RDA2) explained 9.4% of the variance; on the other hand, the two first unconstrained axes (PC1 and PC2) represented less than 8% of the total variance, i.e., much less than that explained by the explanatory variables together.

To uncover other potential underlying mechanisms of mitochondrial regulation, we then sought to examine the relationships existing among all aforementioned data sets by adding a further categorical variable, namely the racing performance of horses. To perform this task, we used the DIABLO framework from mixOmics (Singh *et al*, 2019). While the mt-related genes showed high levels of covariation with the fecal microbiota (*r*^2^ > 0.91, Fig 4A), it was not possible to identify a tight relationship with the other data sets. A more fine-grained view of this biological system was then obtained by focusing on pairwise correlations between variables. The first component of the DIABLO analysis highlighted a significant link between a subset of 45 mt-related genes (all encoded by the nuclear genome) and four gut taxa (i.e., the genus *Mogibacterium,* the species *Eubacterium coprostanoligenes* and the groups *Rikenellaceae* RC9 and *Ruminococcaceae* NK4A214). A link to blood metabolites related to energy supply (i.e., methyl groups of FAs and choline-containing compounds) and metabolites related to the TCA cycle such as glutamine, glutamate, and α-glucose was also unveiled (Fig 4B-C). Moreover, mt-related genes co-occurred with pronounced variations of microbiota derived metabolites, including fecal acetate and valerate, and plasmatic concentrations of TMAO, dimethyl sulfone, and formate. The subset of 45 mt-related genes was functionally enriched in pathways related to fatty acid β-oxidation, mitochondrial apoptosis and biogenesis, respiratory electron transport and signalling and innate immune system response (Table EV12).

**Figure 4.**
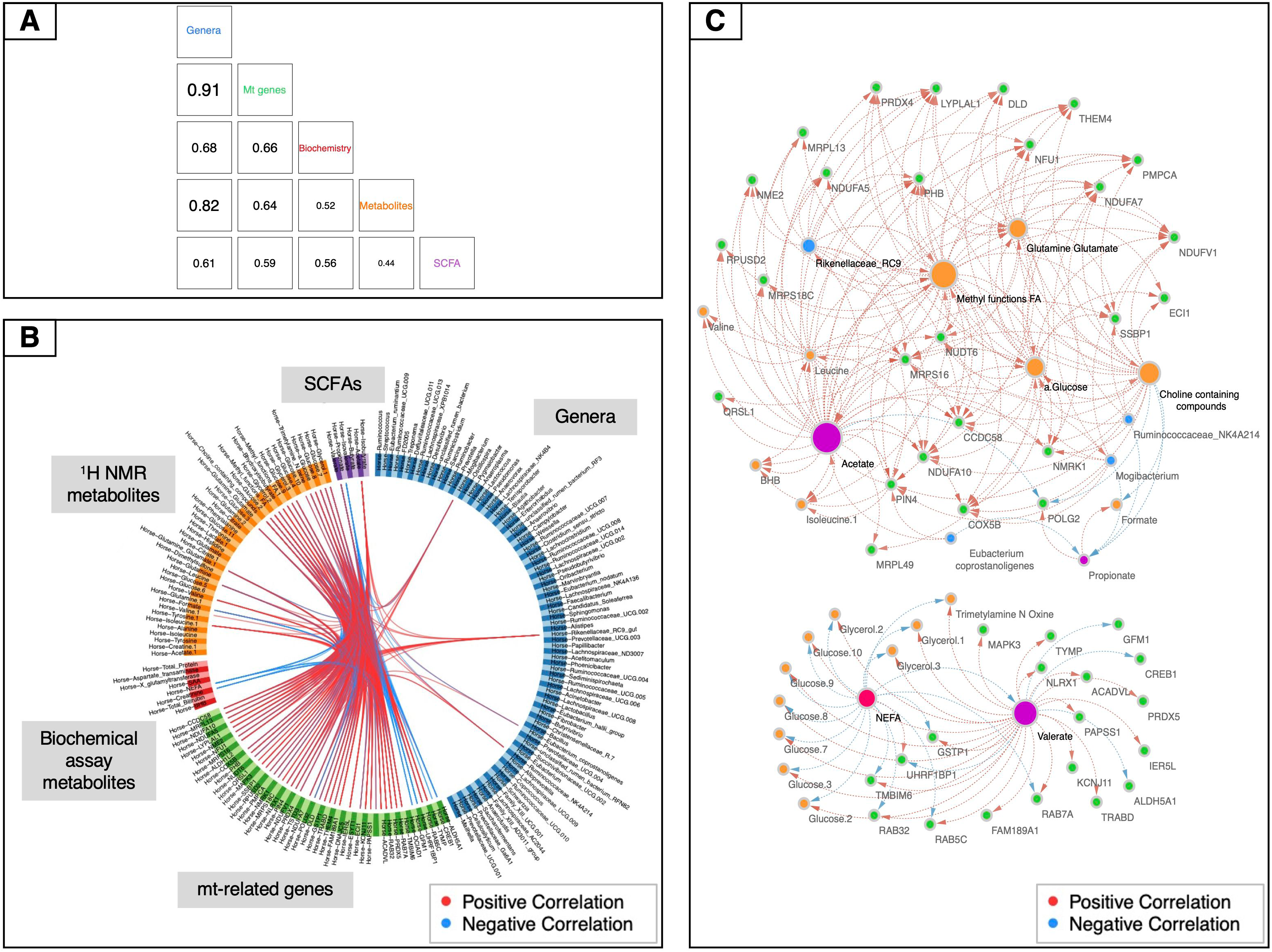
Data integration using mt-related genes, ^1^H NMR metabolites, biochemical assay metabolites, fecal SCFAs and gut bacteria. A Matrix scatterplot showing the correlation between the first components related to each dataset in DIABLO according to the input design. B CIRCOS plot of the final multi-omics final signature. Each dataset is given a different colour: mt-related genes are in green, ^1^H NMR metabolites in orange, biochemical assay metabolites in red, fecal SCFAs in violet and gut bacteria in dark blue. Red and blue lines indicate positive and negative correlations between two variables, respectively (*r* ≥ |0.70|). C Visualization of the network obtained with Cytoscape using the final DIABLO multi-omics signature as an input. Only features with more than 15 connections are shown. The size of the nodes indicates the number of interacting partners within the network.

Finally, an rCCA analysis was also carried out to study in a more targeted way the relationships between mt-related genes and gut microbiota. In this case, the most relevant associations were represented by 90 genes and 9 bacterial genera (Table EV13). Overall, this method largely validated the associations already detected with the other aforementioned approaches. First, all of the four genera highlighted by DIABLO were confirmed, as well as the genus *Fibrobacter*, which had already been detected using NMDS and the db-RDA method. Second, three more taxa found with rCCA appeared to be functionally related to other previously identified microorganisms, thus providing indirect support to those findings. This was the case for *Ruminococcaceae* UCG-002, *Pseudobutyrivibrio* and the *Eubacterium hallii* group.

## DISCUSSION

In this work, we present an integrative study that combines the whole blood transcriptomic with untargeted serum metabolome data, blood biochemical assay profiles and gut metagenome in 20 equine-athletes. We assumed the adaptive response to extreme endurance exercise was essentially explained through the gut-mitochondria axis. Indeed, the different and complementary statistical approaches that we used confirmed this hypothesis, highlighting that the two main omic layers at play were the mt-related genes and the gut microbiota composition (*r*^2^ > 0.91).

Whole blood transcriptome underlines in a clear manner the global response to exercise in equines (Mach *et al*, 2017b, 2016; Capomaccio *et al*, 2013; Barrey *et al*, 2006; Ropka-Molik *et al*, 2017), including the inflammatory response of the muscle associated with sarcolemma permeability and rhabdomyolysis (Barrey *et al*, 2006) (displayed in our study by the high levels of plasma creatine kinase and aspartate aminotransferase after the race). Yet, it remains to be explored whether the whole blood transcriptome reflects the physiological events occurring at the mitochondrial level, notably in the tissues that are highly solicited under endurance, like for instance, the skeletal muscles, the heart and the liver (Gunn, 1987). In addition, whether the transcription of nuclearly-encoded and mitochondrially-encoded genes are regulated in a coordinated way is much less well understood. We therefore compared the transcriptome profile in equine-athletes before and after exercise and we used a meta-analytical approach that allowed us to identify 801 differentially expressed genes that were putatively linked to mitochondria. These genes fit neatly into the well-characterized context of adaptive mitochondrial regulation to endurance and mostly belonged to molecular pathways such as mitochondrial biogenesis, energy metabolism through OXPHOS and FAO, resistance to oxidative stress, mitophagy and inflammation regulation.

We then specifically focused on the mechanisms underlying the biological links between the aforementioned mt-related genes and the gut microbiota to untangle the gut-mitochondria crosstalk. The interdependence of mitochondria and gut microbiota is underscored by several lines of evidences (Yardeni *et al*, 2019; Gruber & Kennedy, 2017; Mottawea *et al*, 2016; Zhang *et al*, 2020; Han *et al*, 2017; Qi & Han, 2018; Ruiz *et al*, 2020; Saint-Georges-Chaumet & Edeas, 2018), although the range and extent of this interplay are largely unknown. For example, (Mottawea *et al*, 2016) showed that butyrate-producing bacteria and mitochondrial proteins were positively correlated, suggesting a signalling role for butyrate in mitochondrial gene expression. In support of this observation, our results revealed that several functionally redundant butyrate-producing bacterial families were associated with the mt-related genes, namely *Lachnospiraceae* (*Oribacterium, Butyrivibrio, Agathobacter* and *Eubacterium* spp*.), Ruminococcaceae*, *Spirochaetaceae* (*Treponema* spp.) (Vital *et al*, 2015; Vacca *et al*, 2020; Gharechahi *et al*, 2020) and *Rikenellaceae* (Vital *et al*, 2015). The bioavailability of butyrate is obviously related to endurance performance because of the role played by this molecule in energy metabolism (Mollica *et al*, 2017). Beyond the scope of its energy producing capacity, butyrate is also known to induce the expression of *PPARγ* gene (Gao *et al*, 2009) and downstream targets in different cells. Our blood transcriptomic analysis indicated an upregulation of *PPARγ* following exercise, raising the possibility that the enrichment in butyrate-producing bacteria increased the expression of this transcription factor and downstream signaling, leading to fatty acid shuttling into and oxidation by the mitochondria. Notably, the redox imbalance during strenuous exercise might be also attenuated by butyrate (Mottawea *et al*, 2016; Dobashi *et al*, 2011). The potential of butyrate to improve exercise capacity has been further posited by Gao *et al*. (2009) and Henagan *et al*. (2015), who observed that the supplementation of this molecule improved the oxidative skeletal muscle phenotype, its mitochondrial content and its proportion of type I fibers. Concomitantly, it is possible that serum lactate, which appeared to be significantly increased in our horses upon prolonged exercise, entered the gut lumen, where it was subsequently transformed into butyrate by *Eubacterium hallii* (Duncan *et al*, 2004; Scheiman *et al*, 2019). Indeed, *Eubacterium hallii* was associated to a subset of mt-related genes according to our rCCA analysis. *Eubacterium*-derived butyrate could then be absorbed into the portal vein and serve as an energy source to the different organs. Another plausible mechanisms employed by microbiota to communicate with mitochondria involved valerate, a branched SCFA formed from protein and amino acid degradation (Fernandes *et al*, 2014). It may be debated whether the urea produced by the host during endurance could be hydrolyzed by commensal gut microbiota resulting in valerate.

Beyond SCFAs, the secondary bile acids could also play an important role in gut-microbiota crosstalk. The genera *Eubacterium* and *Clostridium,* which contributed significantly to our biological system, have the capacity to degrade 5–10% of the primary bile acids forming secondary bile acids (Gérard, 2013). Secondary bile acids might interact with the mitochondria via the activation *FXR-CREB* axis. *CREB*, which was significantly increased in our horses after the endurance race, is a sensor of energy charge and other stress signals, and is a regulator of metabolism that activates autophagy and lipid catabolic functions (Seok *et al*, 2014). Therefore, a picture emerges that, under conditions that foster an increased colonization by these microorganisms, the production of butyrate, valerate and secondary bile acids in the intestine is likely increased, with potential effects on the mitochondria functionality and endurance performance (Fig 5). Yet, evidence of causality of microbiome-derived metabolites on the gut-mitochondria crosstalk remains elusive.

**Figure 5.**
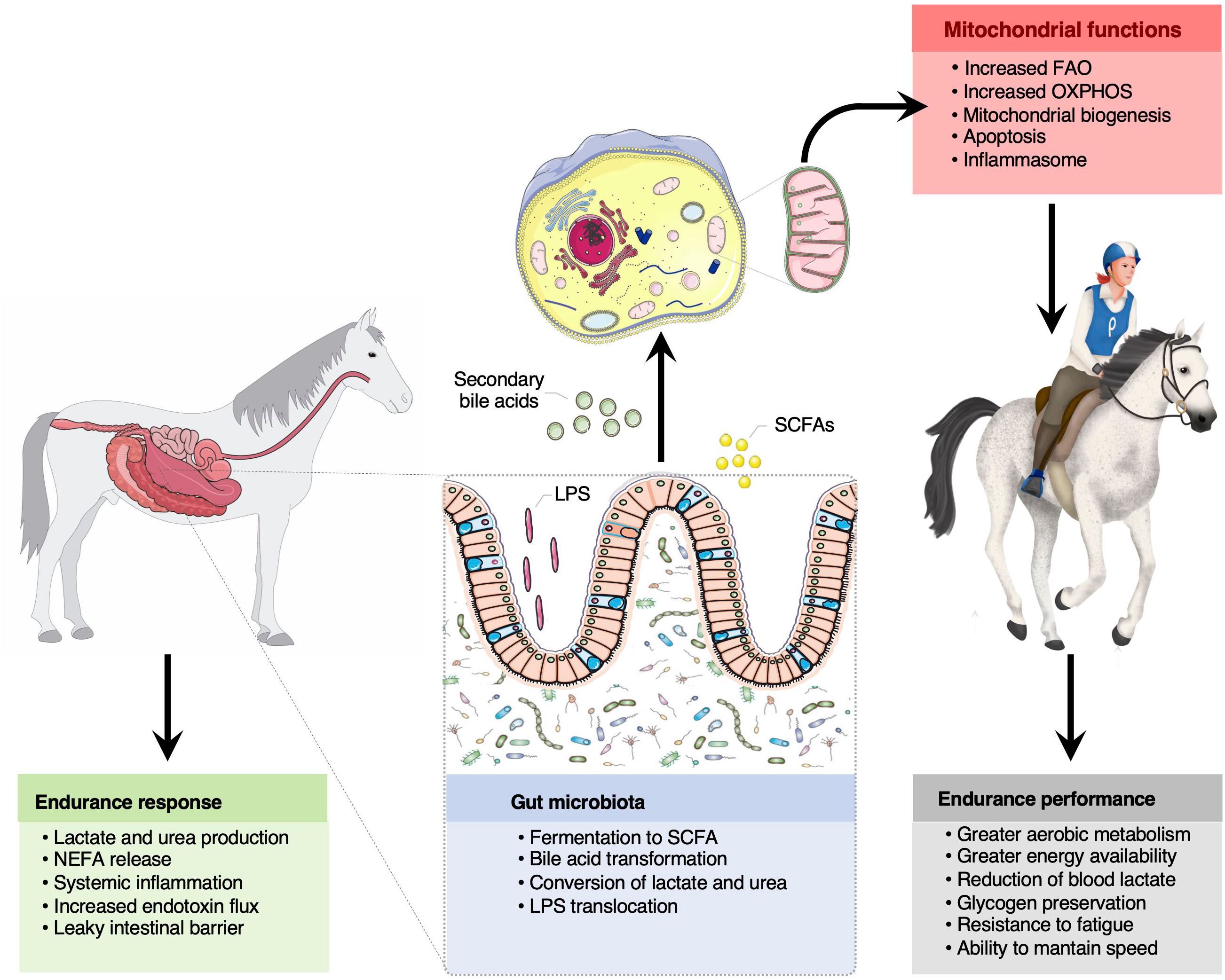
The bidirectional crosstalk between the gut microbiota and mitochondria in endurance horses. The intertwined communication between mitochondria and gut microbiota was likely mediated by microbiota derived byproducts (SCFA and secondary bile acids), which regulate mitochondrial redox balance, inflammation and energy production during intense exercise. Among the SCFA, butyrate appeared as a key regulator of mitochondrial energy production and oxidative stress. Increased lactate and urea concentrations upon prolonged exercise likely entered the gut lumen and were subsequently transformed into SCFA. It is also suggested that circulating free fatty acids participated in the mitochondrial regulation of the inflammatory processes elicited by oxidative stress, microbial dissemination and microbial lipopolysaccharides translocation outside of the gastrointestinal tract, as often occurs in endurance athletes. Whether these mechanisms confer an advantage for endurance performance remains still speculative, but results raise the possibility that gut-microbiota crosstalk is pivotal for greater energy availability, aerobic metabolism, glycogen preservation, resistance to fatigue and to maintain speed during the race. Written permission for publication of the horse drawings was obtained. The pictures of the mitochondria and gut were downloaded from In all cases, no changes were made. Servier Medical Art by Servier is licensed under a <u>Creative Commons Attribution 3.0 Unported License.</u>

The coordination between mitochondria and gut microbiota was presumably regulated by the circulating free fatty acids, the so-called NEFAs. It is becoming clearer that NEFAs not only serves as energy source in the working muscles but act as extracellular signaling molecules that modulate the production of chemokines and cytokines, and the synthesis of pro-inflammatory lipid-derived species (Rodríguez-Carrio *et al*, 2017). Thus, a provocative extension of our work suggests that increased release of NEFAs participated in the mitochondrial regulation of the inflammatory processes elicited by oxidative stress, microbial dissemination and microbial lipopolysaccharides translocation outside of the gastrointestinal tract, commonly observed in endurance athletes (Fielding & Dechant, 2012). Increased release of free fatty acids may dampen the inflammatory response and prevent or mitigate the negative effects of redox imbalance. Supporting this notion, the mitochondrial sirtuin (*SIRT5*) and *SIRT1*, which have an anti-inflammation function (Wang *et al*, 2017), were found to be increased during endurance.

The observations presented herein indicate that the horse could be considered as an interesting *in vivo* model for research in the field of human exercise given its large body size, the aptitude for endurance exercise (Votion *et al*, 2012; van der Kolk *et al*, 2020), that is, high baseline maximal oxygen uptake (VO_2max;_ ∼120 mL·min^-1^·kg^-1^) and the ability to sustain work at a high percentage of VO_2max_ without either the accumulation of exponential levels of blood lactate or skeletal muscle fatigue and the exercise economy (van der Kolk *et al*, 2020; Cottin *et al*, 2010; Goachet & Julliand, 2015). The level of exercise performed by a horse during an endurance competition is similar to that of a human marathon runner (Mach *et al*, 2016; Capomaccio *et al*, 2013) or ultramarathon runner (Scott *et al*, 2009). Nevertheless, despite the usefulness of this model, the differences between the microbiota of horses and humans are relevant. In contrast to what happens in humans, in horses the cecum is large relative to the total gastrointestinal tract and it is an important site for the fermentation of plant materials. Our study presents other limitations. Although the omic approaches used here are considered robust and generate high quality data, they still present several limitations. For instance, 16S rRNA sequencing measures the relative abundance of bacterial genera contained in it, but it does not give any information about its actual functionality, which should be therefore evaluated using other methods, such as for instance metatranscriptomics. Moreover, in our case, ^1^H NMR has been able to detect only metabolites at high concentrations, like in the cases of amino acids, lipids, choline and N-acetylglucosamine. In this regard, the combination of ^1^H NMR and mass spectrometry should result in better coverage of metabolites derived from bacteria, metabolites that are produced by the host and then modified by bacteria and metabolites that are *de novo* synthesized by bacteria. Lastly, it still remains to be determined how the individual components of blood, including plasma, platelets, erythrocytes, nucleated blood cells and exosomes reflect the transcriptomic profiles in horses. Upon endurance, contracting muscles release proteins and metabolites that have endocrine-like properties (Hawley, 2020), but they might also release long non-coding RNAs, myo-miRs and circulating cell-free respiratory competent mitochondria (Al Amir Dache *et al*, 2020; Song *et al*, 2020) that might participate in the aforementioned crosstalk by using the microbiota-derived metabolites.

Taken together, the present study offers extensive novel insight into the mitochondria-gut microbiota axis and opens the way for mechanistic studies that will lead to a better understanding of the orchestrated molecular pathways that underpin endurance adaptations and contribute to the holobiont biology. This is the first description of how metabolites derived from commensal gut microbiota (SCFAs and secondary bile acids), or produced by the host and biochemically modified by gut bacteria (lactate and urea), might influence the genes related to the mitochondria and involved in energy production, redox balance and inflammatory cascades, making them a potential therapeutic target for the endurance. The activation of *PPARγ* and the *FRX-CREB* axis are likely key mechanisms through which SCFAs and bile acids coordinately engage multiple converging pathways to regulate mitochondrial functions, including fatty acid uptake and oxidation to forestall hypoglycemia and ensure longer running time.

Our results also suggest that free fatty acids may not only serve as an important fuel for skeletal muscle during endurance, but may also regulate mitochondrial inflammatory responses through a plethora of mechanisms, the principal one likely being the modulation of the intestinal barrier-ROS production and lipopolysaccharide translocation. Further research focusing on the role that gut microbiota plays on the mitochondrial function across a wide range of tissues and cell types may be highly informative to improve the athlete’s energy metabolism, redox status and inflammatory response.

## MATERIALS AND METHODS

### Ethics approval

The study protocol was reviewed and approved by the local animal care and use committee (ComEth EnvA-Upec-ANSES, reference: 11-0041, dated July 12^th^ 2011) for horse study. All the protocols were conducted in accordance with EEC regulation (n° 2010/63/UE) governing the care and use of laboratory animals, which has been effective in France since the 1^st^ of January 2013. In all cases, the owners and riders provided their informed consent prior to the start of study procedures with the animals.

### Animals

Twenty pure-breed or half-breed Arabian horses (7 females, 3 male, and 10 geldings; mean ± SD age: 10 ± 1.69) were selected from the cohort used by Plancade *et al*. (2019) (Table EV1). The 20 horses were selected following these criteria: (1) enrolment in the 160 km or 120 km category; (2) blood sample collection before and after the race; (3) feces collection before the race; (4) absence of gastrointestinal disorders during the four months prior to enrolment; (5) absence of antibiotic treatment during the four months prior to enrolment and absence of anthelmintic medication within 60 days before the race; (6) a complete questionnaire about diet composition and intake.

Among the 20 horses selected for this study, 16 horses were enrolled for the 160 km category and four for the 120 km category. In the 160 km category, two animals were eliminated due to tiredness after 94 km and 117 km, respectively, and five horses failed a veterinary gate check due to lameness after 94 km (*n* = 1), and after 117 km (*n* = 4). In the 120 km category, one horse was eliminated due to metabolic troubles after 90 km (Table EV1).

The weather conditions, terrain difficulty and altitude were the same for all the participants enrolled in the study as all races (120 and 160 km) took place during October 2015 in Fontainebleau (France). The average air temperature was 15 °C, with a maximum of 20 °C and a minimum of 11 °C, the average air humidity was 88%, and no rain was recorded.

As detailed by Plancade *et al*. (2019), to ensure sample homogeneity, the participating horses were subject to the same management practices throughout the endurance ride and passed the International Equestrian Federation (FEI) compulsory examination before the start. Animals were fed 2–3 hours before the start of the endurance competition with *ad libitum* hay and 1 kg of concentrate pellets. During the endurance competition, all the animals underwent veterinary checks every 20 to 40 km, followed by recovery periods of 40 to 50 minutes (in accordance with the FEI endurance rules). After each veterinary gate check, the animals were provided with *ad libitum* water and hay and a small amount of concentrate pellets.

### Transcriptomic microarray data production and pre-processing

Blood samples for RNA extraction were collected from each animal at T0 and T1 using Tempus Blood RNA tubes (Thermo Fisher). Because blood interacts with every organ and tissue in the body and has crucial roles in immune response, inflammation and physiological homeostasis (Mohr & Liew, 2007), blood-based transcriptome was carried-out as a means for exploring the response to endurance.

Total RNAs were then isolated using the Preserved Blood RNA Purification Kit I (Norgen Biotek Corp., Ontario, Canada), according to the manufacturer’s instructions. RNA purity and concentration were determined using a NanoDrop ND-1000 spectrophotometer (Thermo Fisher) and RNA integrity was assessed using a Bioanalyzer 2100 (Agilent Technologies, Santa Clara, CA, USA). All the 40 RNA samples were processed.

Transcriptome profiling was performed using an Agilent 4X44K horse custom microarray (Agilent Technologies, AMADID 044466). All of the steps were performed by the @BRIDGe facility (INRAE Jouy-en-Josas, France, http://abridge.inra.fr/), as described previously (Mach *et al*, 2017b, 2016).

The horse array was annotated as described by Mach *et al*. (2017b, 2016). In a limited number of cases, a manual annotation step was also included. Probe intensities were background-corrected using the “normexp” method, log_2_ scaled and quantile normalized using the limma package (version 3.1.42.2) (Smyth, 2004) in the R environment (version 3.6.1). Only the probes which presented, on at least two arrays, intensity values at least 10% higher than the 95% percentile of all the negative control probes were kept. Subsequently, controls were discarded and the probes corresponding to genes were summarized. The obtained expression matrix “E1” was processed with the arrayQualityMetrics R package (version 3.42.0) (Kauffmann *et al*, 2009) for quality assessment. No outliers were detected.

### Transcriptome statistical analysis

The differential analysis was performed using the limma R package. A linear model was fitted for each gene, setting the time, the sex, the distance and whether the animal was eliminated from the race as fixed effects, and comparing T1 to T0. The individual was included as a random effect using the “duplicateCorrelation” function (Smyth *et al*, 2005). The *p*-values were Bonferroni corrected setting a threshold of 0.05. The expression matrix “E1” was then used to perform a scaled PCA analysis with FactoMineR R package (version 2.4) (Lê *et al*, 2008).

To confirm the results of the DE analysis, the WGCNA method was also run on the “E1” matrix using the WGCNA R package (version 1.69) (Langfelder & Horvath, 2008). The parameters for the analysis were set as follows: “corFnc” = bicor, “type” = signed hybrid, “beta” = 10, “deepSplit” = 4, “minClusterSize” = 30, and “cutHeight” = 0.1. The eigengenes corresponding to each identified module were correlated individually to all the ^1^H NMR and biochemical assay metabolites, i.e., a set of 56 different molecules (see next paragraphs). A module was then considered positively or negatively associated to this set of molecules if the Pearson *r* correlation values were ≥ |0.65| for at least 5 molecules and if all the corresponding *p*-values were ≤ 1e-05. The positively and the negatively correlated modules defined in this way were merged to obtain a single gene list, which was subsequently compared to the DEG list using a Venn diagram.

Afterwards, a literature-based meta-analytical enrichment test was carried out to assess whether the DEG list and the gene list obtained using WGCNA were enriched in genes related to mitochondria. To this aim, first a consensus list of genes related to these organelles was created (Expanded View Information and Table EV4). This gene list was then annotated in a descriptive way. First, gene symbols were converted into the corresponding KEGG Orthology (KO) codes using the “db2db” tool (https://biodbnet-abcc.ncifcrf.gov/db/db2db.php) from the bioDBnet suite and using the then up-to-date underlying databases. Then, the retrieved KO codes were processed with the “Reconstruct Pathway” tool (https://www.genome.jp/kegg/tool/map_pathway.html) from the KEGG Mapper suite. Eventually, the obtained KEGG pathways underwent some manual editing step to make them easier to interpret, namely (1) only first- and third-level hierarchies including at least 15 genes were kept for visualization; (2) the “Human diseases” first-level hierarchy, with all its child taxonomic terms, was removed; (3) the “Metabolic pathways” third-level hierarchy was discarded as it was redundant with respect to the other third-level hierarchies of the “Metabolism” term (Fig EV4).

Then, the same gene list was intersected with the genes found expressed on the microarray (i.e., the “E1” matrix). The subset thus obtained was separately intersected with the DEG list and the WGCNA gene modules. A Fisher’s exact test and hypergeometric test were then used to evaluate the overrepresentation of genes related to mitochondrial functions in each list.

The intersection between the genes related to mitochondria found on the microarray and the DEGs included 801 genes and was referred to as “mt-related genes”. It was functionally annotated using ClueGO (version 2.5.7) (Bindea *et al*, 2009) by carrying out a right-sided test. Significance was set at a Benjamini-Hochberg adjusted *p*-value of 0.05 and the k-score was fixed at 0.4. Only the “KEGG 30.01.2019” ontology was selected.

### Proton magnetic resonance (^1^H NMR) metabolite analysis in plasma

As described by Plancade *et al*. (2019) and Le Moyec *et al*. (2019) to characterize the metabolic phenotype of endurance horses in detail, we measured ^1^H NMR spectra at 600 MHz for plasma samples. Blood was collected from each horse the day before the event (T0) and within 30 minutes from the end of the endurance race (T1) using sodium fluoride and oxalate tubes in order to inhibit further glycolysis that may increase lactate levels after sampling. All the samples were immediately refrigerated at 4 °C to minimize the metabolic activity of cells and enzymes and to keep metabolite composition as stable as possible, and clotting time was strictly controlled to avoid cell lysis. After clotting, plasma was separated from blood cells and subsequently transported to the laboratory at 4 °C and then frozen at - 80 °C (no more than 5 h later in all cases). Plasma samples were subsequently thawed at room temperature. Using 5 mm NMR tubes, 600 µL of plasma were added to 100 µL deuterium oxide for field locking. The ^1^H NMR spectra were acquired at 500 MHz with an AVANCE III (Bruker, Billerica, MA, USA) equipped with a 5 mm reversed QXI Z-gradient high-resolution probe. Water signal was suppressed with a pre-saturation pulse (3.42 x 10^-5^ W) during a 3s-relaxation delay at the water resonance frequency. The spectrum was divided into 0.001 ppm regions (bins) over which the signals were integrated to obtain intensities. The high- and low-field ends were removed, leaving only the data between 9.5 to 0.0 ppm. The region between 4.5 and 5.0 ppm, which corresponded to the signal of residual water, was also removed. The data were normalized according to the spectra using the probabilistic quotient method (Dieterle *et al*, 2006) and the bins, corresponding to the variables for the statistical analysis, were scaled to unit variance. Further details on sample preparation, data acquisition, data quality control, spectroscopic data pre-processing, and data pre-processing including bin alignment, scaling and normalization are broadly discussed elsewhere (Le Moyec *et al*, 2014).

As specified in Plancade *et al*. (2019), metabolite identification was then performed by using information acquired from other available biochemical databases, namely HMD (http://www.hmdb.ca/), KEGG (https://www.genome.jp/kegg/), METLIN (http://metlin.scripps.edu/), ChEBI (http://www.ebi.ac.uk/Databases/), and LIPID MAPS (http://www.lipidmaps.org/) and the literature (Le Moyec *et al*, 2014; Mach *et al*, 2017b; Le Moyec *et al*, 2019; Jang *et al*, 2017). Each peak was assigned to a metabolite when chemical shifts of peaks in the samples were the same as in the publicly available reference databases or literature (with a shift tolerance level of ± 0.005 ppm), in order to counteract the effects of measurements and pre-processing variability introduced by factors such as pH and solvents. A manual curation for identified compounds was carried out by an expert in horse metabolomics (Le Moyec *et al*, 2014). Eventually, the relative abundance of each metabolite was calculated as the area under the peak (Zheng *et al*, 2011). A total of 50 metabolites was identified, which belonged to the following broad categories: amino acids, including aromatic and branched-chain amino acids, energy metabolism-related metabolites, saccharides, and organic osmolytes (Table EV6). We refer to our previous work (Plancade *et al*, 2019) for more detailed descriptions of the pre-processing and main results of the plasma metabolome data that were used to generate the input files provided with this study.

### Biochemical assay data production

Blood samples for biochemical assays were collected at T0 and T1 using 10 mL BD Vacutainer EDTA tubes (Becton Dickinson, Franklin Lakes, NJ, USA). As detailed in Plancade *et al*. (2019), after clotting the tubes were centrifuged and the harvested serum was stored at 4 °C until analysis (no more than 48 later, in all cases). Sera were assayed for total bilirubin, conjugated bilirubin, total protein, creatinine, CK, β-hydroxybutyrate, ASAT, γ-glutamyltransferase and serum amyloid A levels on a RX Imola analyzer (Randox, Crumlin, UK). The biochemical values obtained are reported in the Table EV7.

### Fecal measurements, 16S data production and analysis

As described by Plancade *et al*. (2019), fresh fecal samples were obtained while monitoring the horses before the race (no more than 24 h before starting the race, in all cases). One fecal sample from each animal was collected off the ground immediately after defecation as described by Mach *et al*. (2017a) and Plancade *et al*. (2019), and three aliquots (200 mg) were prepared. Aliquots for pH determination were kept at room temperature, while aliquots for SCFA analysis and DNA extraction were snap-frozen. Since most of the horses experienced dehydration after the race, the gastrointestinal emptying was significantly delayed and consequently we were not able to recover the feces after the race.

Fecal pH was immediately determined after 10% fecal suspension (wt/vol) in saline solution (0.15 M NaCl solution). SCFAs concentrations were measured as previously described in Mach *et al*. (2017a). The values obtained are described in the Table EV10.

Total DNA was extracted using the EZNA Stool DNA Kit (Omega Bio-Tek, Norcross, Georgia, USA), and following the manufacturer’s instructions. DNA was then quantified using a Qubit and a dsDNA HS assay kit (Thermo Fisher).

The V3-V4 hyper-variable region of the 16S rRNA gene was amplified as previously reported by our team (Mach *et al*, 2017a; Plancade *et al*, 2019; Clark *et al*, 2018; Mach *et al*, 2020; Massacci *et al*, 2019). The concentration of the purified amplicons was measured using a Nanodrop 8000 spectrophotometer (Thermo Fisher) and their quality was checked using DNA 7500 chips onto a Bioanalyzer 2100 (Agilent Technologies). All libraries were pooled at equimolar concentration, and the final pool had a diluted concentration of 5 nM and was used for sequencing. The pooled libraries were mixed with 15% PhiX control according to the protocol provided by Illumina (Illumina, San Diego, CA, USA) and sequenced on a single MiSeq (Illumina, USA) run using a MiSeq Reagent Kit v2 (500 cycles).

The Divisive Amplicon Denoising Algorithm (DADA) was implemented using the DADA2 plug-in for QIIME 2 (version 2019.10) to perform quality filtering and chimera removal and to construct a feature table consisting of read abundance per amplicon sequence variant (ASV) by sample (Callahan *et al*, 2016). DADA2 models the amplicon sequencing error in order to identify unique ASV and infers sample composition more accurately than traditional Operational Taxonomic Unit (OTU) picking methods that identify representative sequences from clusters of sequences based on a % similarity cut-off (Callahan *et al*, 2016). The output of DADA2 was an abundance table, in which each unique sequence was characterized by its abundance in each sample. Taxonomic assignments were given to ASVs by importing SILVA 16S representative sequences and consensus taxonomy (release 132, 99% of identity) to QIIME 2 and classifying representative ASVs using the naive Bayes classifier plug-in (Bokulich *et al*, 2018). The feature table, taxonomy, and phylogenetic tree were then exported from QIIME 2 to the R statistical environment and combined into a phyloseq object (McMurdie & Holmes, 2013). Prevalence filtering was applied to remove ASVs with less than 1% prevalence and in fewer than three individuals, decreasing the possibility of data artifacts affecting the analysis (Callahan *et al*, 2016). To reduce the effects of uncertainty in ASV taxonomic classification, we conducted most of our analysis at the microbial genus level.

The phyloseq (version 1.32.0) (Mcmurdie & Holmes, 2012), vegan (version 2.5.6) (Dixon, 2003) and microbiome packages (version 1.10.0) were used in R (version 4.0.2) for the downstream steps of analysis. The minimum sampling depth in our data set was 10,423 reads per sample. Reads were clustered into 3,385 chimera- and singleton-filtered Amplicon Sequence variants (ASVs) at 99% sequence similarity. ASV counts per sample and ASV taxonomical assignments are available in Table EV8. Data were aggregated at genus, family, order, class and phyla levels throughout the taxonomic-agglomeration method in the phyloseq R package, which merges taxa of the same taxonomic category for a user-specific taxonomic level. The final table obtained after these steps was called “G1” and included 100 genera (Table EV9).

### qPCR quantification of bacterial, fungal and protozoan concentration

As detailed by Plancade *et al*. (2019), concentrations of bacteria, anaerobic fungi and protozoa in fecal samples were quantified by qPCR using a QuantStudio 12K Flex platform (Thermo Fisher Scientific, Waltham, USA). Primers for real-time amplification of bacteria (FOR: 5’-CAGCMGCCGCGGTAANWC-3’; REV: 5’-CCGTCAATTCMTTTRAGTTT-3’), anaerobic fungi (FOR: 5’-TCCTACCCTTTGTGAATTTG-3’; REV: 5’-CTGCGTTCTTCATCGTTGCG-3’) and protozoa (FOR: 5’-GCTTTCGWTGGTAGTGTATT-3’; REV: 5’-CTTGCCCTCYAATCGTWCT-3’), are described in Mach *et al*. (2015) and Clark *et al*. (2018) and were purchased from Eurofins Genomics (Ebersberg, Germany).

Amplified fragments of the target amplicons were used to create a seven-point 10-fold standard dilution series. The dilution points ranged from 2.25 × 10^7^ to 2.25 × 10^13^ copies per μg of DNA for bacteria and protozoa and from 3.70 × 10^6^ to 3.70 × 10^12^ copies per μg of DNA for anaerobic fungi. qPCR reactions were performed in a final volume of 20 μL, containing 10 μL of Power SYBR Green PCR Master Mix (Thermo Fisher), 2 μL of standard or DNA template at 0.5 ng/μL and 0.6 μM of each primer to a final concentration of 200 mM for bacteria and anaerobic fungi and 150 mM for protozoa.

In all the cases, the thermal cycling conditions were as follows: initial denaturation at 95 °C for 10 min; 40 cycles of denaturation at 95 °C for 15 sec, annealing and extension at 60 °C for 60 sec. To check for the absence of nonspecific signals, a dissociation step was added after each amplification. It was carried out by ramping the temperature from 60 °C to 95 °C. All qPCR runs were performed in triplicate, and the standard curve obtained using the target amplicons was used to calculate the number of copies of microorganisms in feces.

Taking into account the molecular mass of nucleotides and the amplicon length, the number of copies was obtained using the following equation: copies per nanogram = (NL × A × 10^-9^)/ (n × mw), where “NL” is the Avogadro constant (6.02 x 10^23^ molecules per mole), “A” is the molecular weight of DNA molecules (ng), “n” is the length of the amplicon in base pairs, and “mw” is the molecular weight per base pair. The final values obtained are described in the Table EV11.

### Integrative statistical analyses

Data integration was carried out using several approaches and different combinations of data sets. Prior to integration, each data set underwent a specific set of pre-processing steps. In the case of the ^1^H NMR and biochemical assays, data were processed by subtracting the T0 values from the T1 values. The two data sets included 50, and 9 molecules, respectively.

In the case of the blood transcriptome, a new expression matrix (“E2”) was created including only the differentially expressed mt-related genes and by subtracting the T0 from the T1 expression matrix values, i.e., by calculating the ratio between T1 and T0 log scaled expression values from the two matrices. A total of 801 genes was retained (Table EV5).

The fecal pH values and the proportions of the 6 SCFAs in feces did not undergo any specific pre-processing. In the case of fecal microbiota, the genera table “G1” was modified using the mixMC framework of the mixOmics R package (version 6.10.9) (Rohart *et al*, 2017). First, raw data were pre-filtered by removing the genera for which the percentage of the sum of counts was lower than 1% compared to the total sum of all counts. Then, the pre-filtered data were transformed using the Centered Log Ratio transformation (CLR) and applying an offset of 1. The filtered genera matrix “G2” obtained in this way included 85 genera (Table EV14). Finally, the concentration of microorganisms in feces did not undergo any specific editing.

After the pre-processing steps, a first round of integration was performed using three different methods and working with all six data sets available, namely: (1) mt-related genes (i.e., the “E2” matrix); (2) ^1^H NMR metabolites; (3) biochemical assay metabolites; (4) the concentrations of fecal SCFAs; (5) fecal 16S rRNA gene sequencing data (i.e., the “G2” matrix); and (6) the concentration of fecal microorganisms.

As a first integration approach, a global NMDS ordination was used to extract and summarize the variation in mt-related genes (the “response variable”) using the “metaMDS” function in vegan R package. To determine the number of dimensions for each NMDS, stress values were calculated. Stress values are a measure of how much the distances in the reduced ordination space depart from the distances in the original p-dimensional space. High stress values indicate a greater possibility that the structuring of observations in the ordination space is entirely unrelated to that of the original full-dimensional space.

The other five data sets (the “explanatory variables”) were then fitted to the ordination plots using the “envfit” function in the vegan R package (Clarke & Ainsworth, 1993) with 10,000 permutations. The “envfit” function performs a multivariate analysis of variance (MANOVA) and linear correlations for categorical and continuous variables. The effect size and significance of each covariate were determined comparing the difference in the centroids of each group relative to the total variation, and all of the *p*-values derived from the “envfit” function were Benjamini-Hochberg adjusted. The obtained *r*^2^ gives the proportion of variability (that is, the main dimensions of the ordination) that can be attributed to the explanatory variables.

As a second integration approach, a forward-selection model-building method for redundancy analysis (RDA) (Blanchet *et al*, 2008) was used to extract and summarize the variation in mt-related genes (the “response variable”) that could be explained by the other five data sets (the “explanatory variables”). To determine which set of covariates provided the most parsimonious model, automatic stepwise model selection for constrained ordination methods was used as implemented by the “ordistep” function of the vegan R package. To test for robustness, a forward automatic model selection on a distance based RDA was then performed using the “ordiR2step” function of the vegan package in R (Oksanen *et al*, 2013). This provided an estimation of the linear cumulative effect size of all the identified non-redundant covariates and of their independent fraction in the best model. In the case of this latter function, the “E2” matrix was modified using the Hellinger transformation prior to the analysis. These two RDA functions use different criteria for variable selection. The “ordistep” funciton uses the Akaike’s information criterion (AIC) and *p*-value < 0.05 obtained from Monte Carlo permutation tests, while “ordiR2step” uses the adjusted coefficient of determination (*r*^2^_adj_). In both cases, the procedure begins by comparing a null model containing no variables and a test model containing one variable, where every possible covariate is considered.

As a third integrative approach, the N-integration algorithm DIABLO of the mixOmics R package was used. In this case, the relationships existing among all six data sets were studied by adding a further categorical variable, i.e., the performance of horses. Horses that had a poor performance or that had been eliminated (*n* = 8) were compared to horses that had completed the race (*n* = 12, Table EV1). DIABLO seeks to estimate latent components by modelling and maximizing the correlation between pairs of pre-specified datasets to unravel similar functional relationships between them (Singh *et al*, 2019). A full weighted design was considered and, to predict the number of latent components and the number of discriminants, the “block.splsda” function was used. In both cases, the model was first fine-tuned using the leave-one-out cross-validation by splitting the data into training and testing. Then, classification error rates were calculated using balanced error rates (BERs) between the predicted latent variables with the centroid of the class labels (i.e., eliminated vs non eliminated horses) using the “max.dist” function. BERs account for differences in the number of samples between different categories. Only interactions with *r* ≥ |0.70| were visualized using CIRCOS. To visualize the high-confidence molecule co-associations determined by CIRCOS, only those with *r* ≥ |0.70| and more than 15 connections were automatically visualized using the organic layout algorithm in Cytoscape (version 3.8.1) (http://cytoscape.org).

Finally, we performed a pairwise integration, focusing only on mt-related genes and microbiota data, and using the rCCA method as implemented by the mixOmics R package (https://cran.r-project.org/web/packages/mixOmics/). The penalization parameters were estimated using the “shrinkage” method and setting the “ncomp” parameter to two. The correlation matrix thus obtained was filtered by retaining only the genes and the genera for which at least one association value data point presented *r* ≥ |0.55|.

## ACKNOWLEDGMENTS

We are grateful to Marine Beinat, Julie Rivière, Emmanuelle Rebours, Jordi Estellé, Caroline Morgenthaler and Arni Janssen for participating in the sample collection and organization during the project. We particularly thank Valentine Ballan and Fanny Blanc who helped to isolate the blood RNA. We also thank Catherine Philippe for the SCFA analysis in feces, Diane Esquerré who prepared the libraries and performed the MiSeq sequencing at the GeT-PlaGe genomics core facility (INRAE, Toulouse, https://get.genotoul.fr/) and Estel Blasi, who created the picture of horses. Lastly, we are grateful to the INRAE MIGALE bioinformatics facility (MIGALE, INRAE, 2020, Migale bioinformatics facility, doi: 10.15454/1.5572390655343293E12) for providing computing and storage resources and to all the horse owners, riders and endurance race organizers who participated in the study. This work was funded by grants from the *Fonds Eperon*, the *Institut Français du Cheval et de l’Equitation* (IFCE), the *Association du Cheval Arabe* (ACA) and the *Institute National de la Recherche pour l’Agriculture, l’alimentation et l’environnement* (INRAE).

## AUTHOR CONTRIBUTIONS

NM and EB conceived of the presented idea. NM designed the experiment, performed the PCR and the RT-qPCR experiments, carried out most of the bioinformatics analysis and the integrative biology approaches. NM wrote the main manuscript text and prepared the figures with support from MM and EB. MM carried out the transcriptome analyses, the rCCA integrative analysis and prepared most of the tables. MM and EB created the consensus list of genes related to mitochondria. AR verified the statistical methods and CR verified the blood biochemical and the performance data. JL performed all the laboratory steps related to microarray analyses. LLM performed the metabolomic experiment and analyzed the metabolite peaks. CR and EB were in charge of the organization and sampling management during the race. All authors reviewed the manuscript and approved the final version.

## CONFLICT OF INTEREST

The authors declare no competing interests.

## DATA AVAILABILITY SECTION

The datasets produced in this study are available in the following databases:

- Microarray expression data: Gene Expression Omnibus (GEO) repository under the accession number GSE163767; (https://www.ncbi.nlm.nih.gov/geo/query/acc.cgi?acc=GSE163767)
- Metabolomic data: NIH Common Fund’s Data Repository and Coordinating Center UrqK1489; (http://dev.metabolomicsworkbench.org:22222/data/DRCCMetadata.php?Mode=Study&StudyID=ST000945)
- Gut metagenome 16S rRNA targeted locus data: DDBJ/EMBL/GenBank under the accession KB000000, version KB000000.1; (locus KB000000). The corresponding BioProject is PRJNA438436, and the accession numbers of the BioSamples included in it were SAMN08715709 to SAMN08715760.

## EXPANDED VIEW INFORMATION

### Creation of a consensus list for literature-based meta-analytical analysis of mitochondrial-related genes

We retrieved four gene lists: (1) the Integrated Mitochondrial Protein Index (IMPI) gene list (Smith & Robinson, 2019). Only the human gene list (‘Human IMPI genes’, ‘MitoMiner version Q2 2018’, downloaded from http://mitominer.mrc-mbu.cam.ac.uk/release-4.0/impi.do) was retained (1,626 genes) to facilitate finding horse/human orthologs; (2) the Mitocarta Inventory (Calvo *et al*, 2016) gene list. As before, only the human gene list (‘Human MitoCarta 2.0 genes’, ‘MitoCarta 2.0 genes in MitoMiner’ downloaded from http://mitominer.mrc-mbu.cam.ac.uk/release-4.0/mitocarta.do) was retained (1,158 genes) to facilitate the identification of horse/human orthologs; (3) all of the 618 genes included in the following KEGG pathways: hsa00020, hsa00061, hsa00062, hsa00071, hsa00072, hsa00100, hsa00130, hsa00140, hsa00190, hsa00240, hsa00280, hsa00290, hsa00471, hsa00480, hsa00628, hsa00760, hsa00860, hsa01212, hsa04020, hsa04024, hsa04072, hsa04115, hsa04136, hsa04137, hsa04140, hsa04210, hsa04215, hsa04350, hsa04370, hsa04668 and hsa04979. The genes in each KEGG pathway were retrieved from the KEGG database (release 96.0) using the R package KEGGREST; (4) a custom list including 103 genes found in the literature (Pearce *et al*, 2017; Bianchessi *et al*, 2016; Nicholls & Gustafsson, 2018; Wang *et al*, 2016; Cosson *et al*, 2012; Rizzuto *et al*, 2012; Gustafsson & Samuelsson, 2001; Gustafsson *et al*, 2016; Lee *et al*, 2015). These four gene sets were merged to create a consensus list, which included 2,082 unique genes (Table EV4).

## EXPANDED VIEW FIGURES

**Figure EV1.**
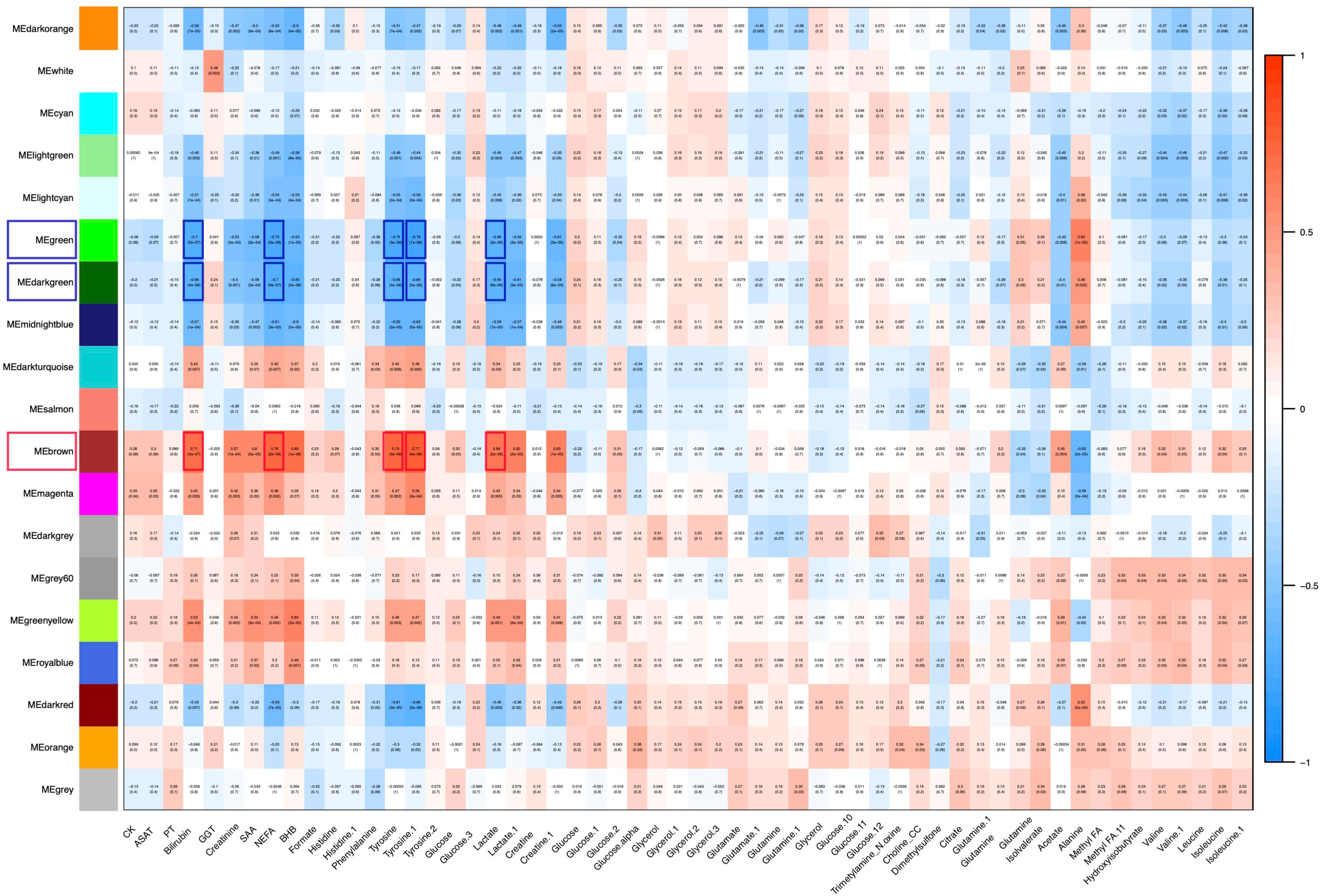
Correlations between eigengene modules and metabolites from WGCNA method. Each module is labelled with a unique colour as an identifier. A module is considered positively (highlighted in red) or negatively (highlighted in blue) associated to metabolites if the Pearson *r* correlation values were ≥ |0.65| for at least 5 molecules (highlighted again in red or blue) and if the corresponding *p*-values are ≤ 1e-05. Within each cell, upper values indicate the correlation values between modules and metabolites, while lower values are the corresponding *p*-values.

**Figure EV2.**
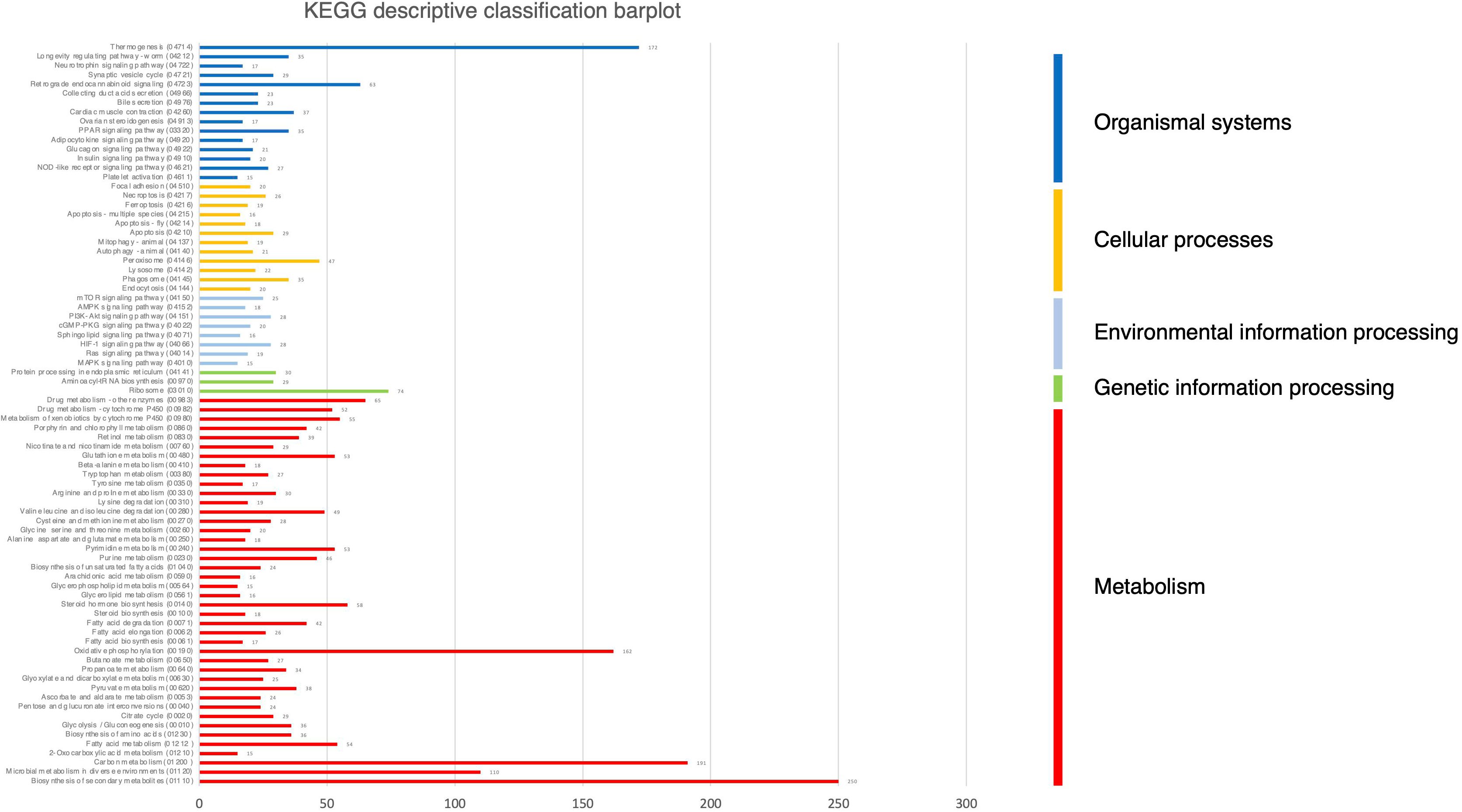
Descriptive KEGG pathway classification bar plot obtained using the consensus list of genes related to mitochondria. The horizontal bars represent the absolute number of genes found in third-level KEGG pathways, grouped in first-level KEGG pathways using a colour code. The vertical bars on the right indicate the names of first-level pathways.

**Figure EV3.**
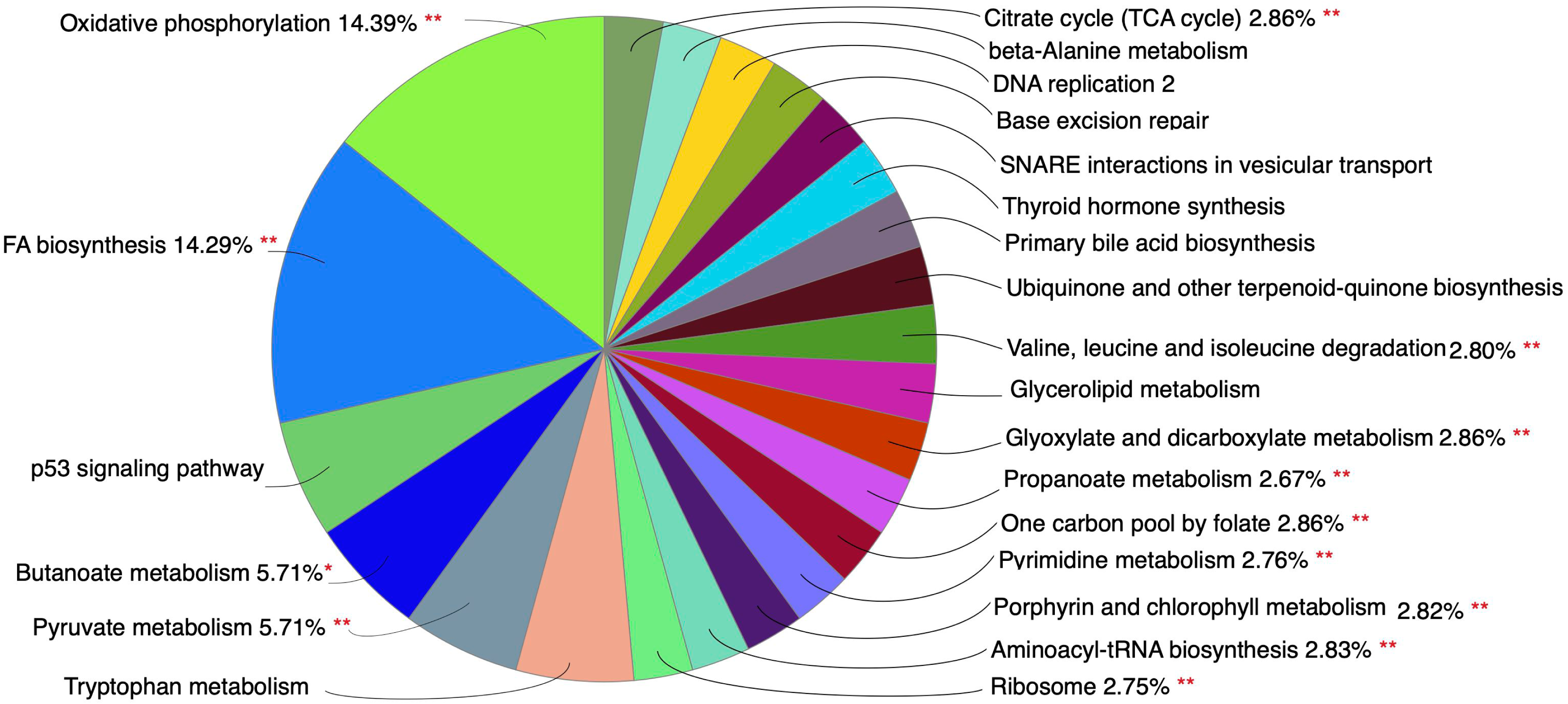
Functional classification of the mt-related genes. Functional classification of mt-related genes obtained with ClueGO. The chart shows the functional groups found as enriched, and the name of the group is represented by the group leading term. The significant enriched functional groups are marked with a red *.

**Figure EV4.**
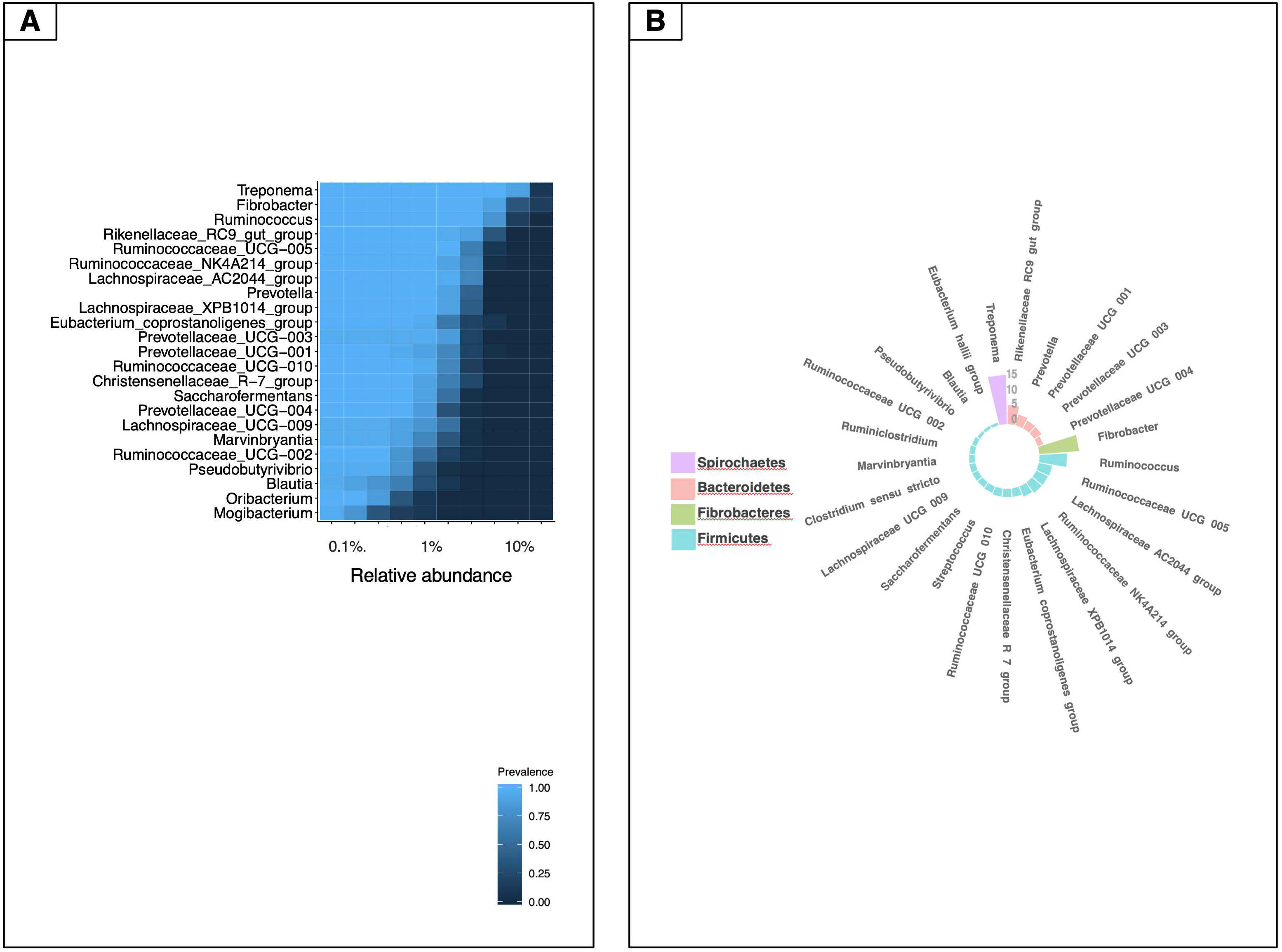
Composition of the core microbiota. A heatmap showing the core microbiota and its prevalence at different detection thresholds. Only the genera shared by 99% of individuals in the cohort and with a minimum detection threshold of 0.1% are shown. B Circular stacked barplot of the main genera included in the core genome. Genera are coloured according to the phylum they belong to.

## EXPANDED VIEW LARGE TABLES

**Table EV1 - Metadata of the horses recruited in the experiment.**

The letters between brackets in the “Ranking” column indicate the causes of the elimination of the horse: “L” corresponds to “lameness”, R” to “retired” and “M” to “metabolic issues”.

**Table EV2 - List of genes found differentially expressed in whole blood in T1 with respect to T0 horses.**

Columns two and three show the mean log_2_ normalized expression values in the two time points, while columns four and five show the log_2_ FC values and the Bonferroni adjusted *p*-values. Columns six to nine indicate the genes that are also found in the mt-related consensus list. The gene lists used to create the final consensus are indicated separately.

**Table EV3 - List of genes included in each of the three eigengene modules found as correlated to metabolites using WGCNA.**

The modules are indicated using the default WGCNA naming convention, and highlighted in blue to indicate negative correlation and red to indicate positive correlation.

**Table EV4 - Consensus list of genes related to mitochondria obtained as described in the Expanded View Information.**

Gene description, gene localizations and gene types were determined according to IPA (https://www.qiagenbioinformatics.com/products/ingenuity-pathway-analysis).

**Table EV5 - List of mt-related genes, defined as the intersection between the list included in the Table EV4 and the set of expressed genes.**

Gene description, gene localizations and gene types were determined according to IPA (https://www.qiagenbioinformatics.com/products/ingenuity-pathway-analysis). Columns from N from AG show the expression values found in the “E2” matrix, which were obtained by subtracting the T0 from the T1 expression matrix values, i.e., by calculating the ratio between T1 and T0 log scaled expression values from the two matrices.

**Table EV6 - Relative abundance of metabolites obtained from blood of the 20 horses under study collected before and after the endurance ride.**

**Table EV7 - Biochemical parameters obtained from the blood of the 20 horses under study collected before and after the endurance race.**

**Table EV8 - ASV taxonomical assignments and ASV counts for the 20 horses under study.**

**Table EV9 - Non-normalized annotated abundance genera table observed in the fecal samples of the 20 horses under study.**

**Table EV10 - Fecal pH and fecal short chain fatty acids measurements in the 20 horses under study before the endurance race.**

**Table EV11 - Concentrations of bacteria, ciliate protozoa and anaerobic fungi in the feces of the 20 horses under study.**

**Table EV12 - Correlated variables obtained using DIABLO on all of the available data sets.**

Gene descriptions and localizations were determined using the IPA database (https://www.qiagenbioinformatics.com/products/ingenuity-pathway-analysis). Molecular pathways were determined using ClueGO 2.5.7 in the case of genes.

**Table EV13 - Correlation matrix of the associations between mt-related genes and bacterial genera obtained using the rCCA method.**

Only the genes and the genera for which at least one association value data point presented *r* ≥ |0.55| are shown.

**Table EV14 - Genera table obtained using the mixMC framework.**

Genera with less than 1% counts with respect to the total number were removed, and subsequently a centered log ratio transformation was applied.

